# Pharmacological inhibition of the ALK axis elicits therapeutic potential in Consensus Molecular Subtype 1 colon cancer patients

**DOI:** 10.1101/2020.10.07.307991

**Authors:** Martina Mazzeschi, Michela Sgarzi, Donatella Romaniello, Valerio Gelfo, Carola Cavallo, Spartaco Santi, Michelangelo Fiorentino, Gabriele D’Uva, Balázs Győrffy, Ruth Palmer, Mattia Lauriola

## Abstract

In the last years, several efforts have been made to classify colorectal cancer (CRC) into well-defined molecular subgroups, representing the intrinsic inter-patient heterogeneity, known as Consensus Molecular Subtypes (CMSs). In this work, we performed a meta-analysis of 1700 CRC patients stratified into four CMSs. We identified a negative correlation between a high level of anaplastic lymphoma kinase (ALK) expression and relapse-free survival, exclusively in CMS1 subtype. Stemming from this observation, we tested several CMSs *in vitro* models with crizotinib (CZB) or alectinib (ALC), potent ALK inhibitors, already approved for clinical use. ALK interception strongly inhibits cell proliferation already at nanomolar doses, specifically in CMS1 cell lines, while no effect was found in CMS2/3/4 groups. Furthermore, *in vivo* imaging identified a role for ALK in the dynamic formation of 3D spheroids, which was impaired by the pharmacological inhibition of ALK. Consistently, CZB was responsible for the dampened activation of ALK along with the downstream AKT cascade. Mechanistically, we found a specific pro-apoptotic effect of ALK inhibition in CMS1 cell lines, both in 2D and 3D. Confocal analysis suggests that inhibition in CMS1 cells enhances cell-cell adhesion when growing in 3D. In agreement with our findings, an ALK signature encompassing 65 genes statistically associated with worse relapse-free survival in CMS1 subtype. Finally, the efficacy of ALK inhibition treatment was demonstrated in patient-derived organoids. Collectively, our findings suggest that ALK inhibition may represent an attractive therapy for CRC, and CMS classification may provide a useful tool to identify patients who could benefit from this treatment. These findings offer rationale and pharmacological strategies for the treatment of CMS1 CRC.

## Introduction

Several efforts have been made to stratify colorectal cancer patients into molecularly homogeneous subgroups, in order to unveil prognostic and predictive factors, which may identify specific treatment regimens. To assemble and organize CRC classifications, obtained by different approaches (1–5), Guinney and colleagues developed a unifying association network that successfully identified four consensus molecular subtypes (CMSs), each one with specific biological features (6). Notably, these transcriptional subtypes are maintained in CRC cell models (3,7), thus allowing the selection of a precise *in vitro* model to recapitulate at the best the investigated patient subtypes.

Anaplastic lymphoma kinase (ALK) is a large, glycosylated receptor tyrosine kinase (RTK) that belongs to the superfamily of the insulin receptors, showing a great similarity to leukocyte receptor tyrosine kinase (LTK, 79% identity) (8,9). RTKs undergo ligand-mediated activation, resulting in receptor phosphorylation in response to ligand (10–12). The small secreted ALKAL proteins (ALKAL1 and ALKAL2), also known as FAM150A/B and augmentor-α/β, have been reported as activating ligands for ALK (13,14). Physiologically ALK is expressed in the nervous system to regulate cells proliferation, differentiation and survival, and its expression levels decrease after birth (15). ALK is generally poorly expressed in normal adult tissues, making it a highly promising molecular target for cancer therapy (16). Indeed, ALK is known to be oncogenic in different types of cancer such as Non-Small Cell Lung Cancer (NSCLC) and it has a critical role in neuroblastoma (17), both as a consequence of gene translocation (18), overexpression, point mutations (11,19) or hyper-phosphorylation of the downstream pathways (20). ALK gene copy number gain was also detected in both the alveolar (88%) and embryonal (52%) rhabdomyosarcoma subtypes (21). The involvement of the ALK pathway in colorectal cancer has so far been poorly investigated. CRC patients harbouring *ALK* gene amplification or copy number gain display worse prognosis (22,23), while *ALK* genomic rearrangements, such as translocations or fusion proteins, are rarely identified (7,24-26).

Here we show that *ALK* expression has a robust predictive ability in the survival of CRC patients belonging to CMS1. Stemming from these findings, we developed *in vitro* molecular assays, probing the inhibition of ALK to investigate the benefit of therapies targeting this pathway. Mechanistically, we observed activation of ALK signaling by ALKAL ligands that was blocked by ALK inhibitors in CRC cell lines. Furthermore, these results were coupled with confocal imaging of the 3D spheroids growth. Strikingly, ALK inhibition was effective in terms of proliferation and signalling in CMS1, while no effect was detected in the other subtypes. We also confirmed that 65 genes belonging to the ALK signature were associated with survival specifically in the CMS1 dataset. Finally, evaluation of patients-derived organoids confirmed the therapeutic implication of our findings. In conclusion, our data propose the ALK pathway as a therapeutic target in CMS1 colorectal cancer patients.

## RESULTS

### *ALK* is predictive of survival in consensus molecular subtype 1 CRC patients

Various colorectal cancer patients display a different set of genomic alterations that lead to a diverse response to therapies. To investigate ALK involvement in colorectal cancer, we started from the analysis of a dataset of approximately 1700 CRC patients, stratified for consensus molecular subtypes: CMS1, CMS2, CMS3 and CMS4 (6,27). Patients were tested for the amount (high/low) of mRNA expression of ALK (probe 208212_s_at). The relapse-free survival (RFS) probability of these subjects was calculated throughout 200 months (about 16 years). After trichotomization of the dataset, in each molecular subtype separately, the median group was excluded and the groups with high and low expression of ALK mRNA were compared. High levels of ALK were predictive of poor relapse free-survival specifically in CMS1, with an impressive HR of 2.77 and a P value of 0.0072 (Fig.1). Indeed, most of these patients die within 50 months, suggesting that ALK plays a significant role in CMS1. This trend was specific for CMS1, as CMS2, 3 and 4 subgroups displayed no survival predictive role for ALK (Fig.1). CMS1 is characterized by microsatellite instability, and it is linked to hypermutation and hypermethylation (CIMP). It represents 15% of CRCs, which are triggered by alterations in the DNA mismatch-repair system, inducing microsatellite instability (MSI) and a high mutation rate (28). Next, we searched for a suitable *in vitro* model recapitulating the features of CMS1 subtype. In the literature, some reports classify CRC cell lines according to the CMS, following the same classification system established for patients (7). We employed the classification from Berg and colleagues (29), that assigned each line to its specific CMS, corresponding to the patients’ classification. We selected LoVo and SW48 cells as representative models for CMS1 (where ALK is predictive of poor survival), HT29 and LS174T for CMS3 and HCT116 and Caco-2 for CMS4. We confirmed the lack of ALK aberrant translocations in these cell lines by FISH staining (Supplementary Fig. 1).

**FIGURE 1.**
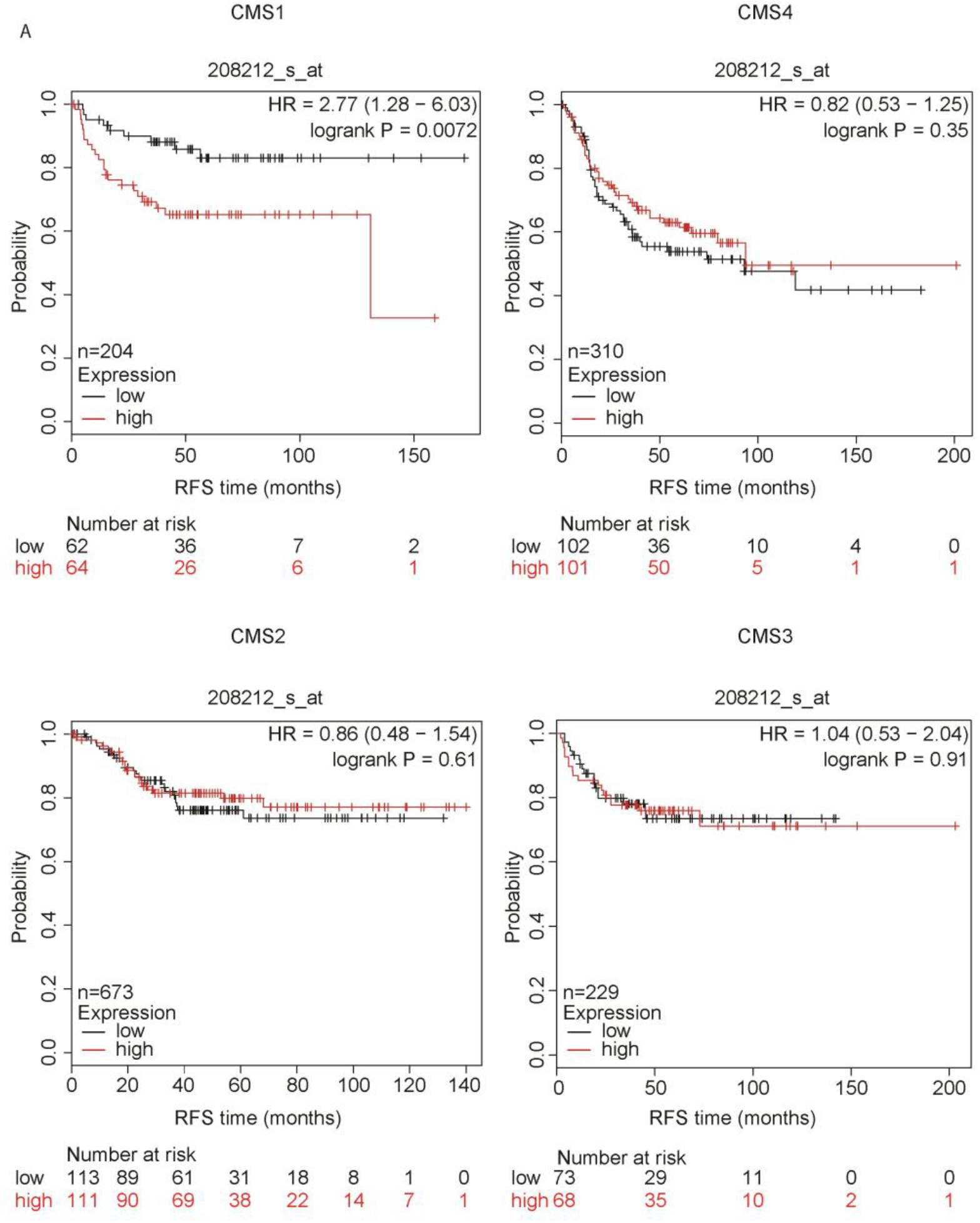
ALK IS PREDICTIVE OF SURVIVAL IN CONSENSUS MOLECULAR SUBTYPE 1. Kaplan-Meier analysis of a cohort of 1700 patients stratified by CRC classification and *ALK* gene expression (probe 208212_s_at). The red line represents high level of expression, while the black one is representative of low levels. For each patient, the relapse-free survival (RFS) probability is reported over a period of maximum 200 months. Notably, a higher expression of ALK mRNA is predictive of a worse RFS only in consensus molecular subtype 1 (CMS1), with an HR of 2.77 and p-value: 0.0072. No statistically significant association was observed between ALK expression and RFS in the other subgroups.

### Antitumorigenic action of ALK axis inhibition is specific for CMS1

We employed two different ALK inhibitors, crizotinib (CZB) and alectinib (ALC), which are currently used for the treatment of ALK-positive NSCLC patients. To experimentally examine the patients data-driven hypothesis of ALK pathway involvement in CMS1 colon cancer subtype, we treated CMS1 (LoVo and SW48), CMS3 (HT29 and LS174T) and CMS4 (HCT116 and Caco-2) cell lines with increasing concentrations of CZB (Fig. 2A) and ALC (Supplementary Fig. 2). ALK inhibition strongly affects LoVo and SW48 cells proliferation. Indeed, CZB inhibited proliferation with a half maximal inhibitory concentration (IC50) of about 9.6nM and 256nM for LoVo and SW48 respectively, while almost no effect was recorded in the other cell lines (Fig. 2A). These results were consistent with the colony forming ability assay, that measures the percentage of cells retaining the capacity of producing several progenies, detected as colonies. Cells seeded at very low density were treated in triplicate with increasing concentrations of CZB or ALC. After ten days and before reaching a saturating full confluence, cells were fixed and stained (Fig. 2B and Supplementary Fig. 2A). Notably, at doses of 250 nM, very few colonies were detected in LoVo and SW48 cells, while minimal effect was observed in HCT116 and Caco-2 cells or LS174T and HT-29 (Fig. 2B and Supplementary Fig. 2A). Consistently, in CMS1 cell lines, the S phase entry evaluation by BrdU incorporation was strongly suppressed (Fig. 2C). Altogether, these data suggest that CMS1 cell lines depend on ALK pathway for proliferation and survival, while CMS3 and 4 cells are less vulnerable to its inhibition. To further validate these results, we tested ALK inhibition by CZB and ALC, on a system closer to tumours *in vivo*. Cells were seeded in adhesion lacking conditions, forced to grow in three-dimensional structures, known as spheroids, as previously reported (30,31). Multicellular spheroids reproduce the 3D architecture, cell-cell interactions as well as oxygen and proliferation gradients observed in tumours (32). Spheroids were left to grow for approximately 15 days and the size of each spheroid under the vehicle or CZB/ALC treatment was measured. In these conditions, we found that ALK inhibition strongly affects LoVo and SW48 growth (Supplementary Fig. 3A and 2B). On the other hand, HCT116 and HT-29 cells cultured in the same conditions performed very well, growing completely undisturbed, even at high doses of drugs (Supplementary Fig. 3A), suggesting that ALK inhibition is not active on CMS3 and CMS4 cells, while CMS1 are highly sensitive. Next, we applied a soft agar assay, which represents a surrogate assay for tumorigenesis. It assesses the impact of inhibitors on cancer cell 3D colony formation and is frequently the first experimental validation that compounds, already known to be active in 2D, must possess to progress through the drug discovery pipeline. Under these growing conditions, ALK inhibition of proliferation resulted again impressive in CMS1 cell lines, while no effect was detected in CMS4 cells (Supplementary Fig. 3B and 2C). Previous studies reported that CZB induces apoptosis in colorectal cancer cell lines, in a process involving upregulation of PUMA (33). Thus, we evaluated apoptosis at single cell level, by the terminal deoxynucleotidyl transferase (TdT)-mediated deoxyuridine triphosphate (dUTP)-biotin nick end-labelling (TUNEL) method. ALK inhibition during a timeframe of 48h was sufficient to increase TUNEL positivity in CMS1 cells (Fig. 3A). Consistently, the mRNA levels of the antiapoptotic marker BCL2 decreased, while the pro-apoptotic BAX steadily increased, reaching the maximum at 48h (Fig. 3B).

**FIGURE 2.**
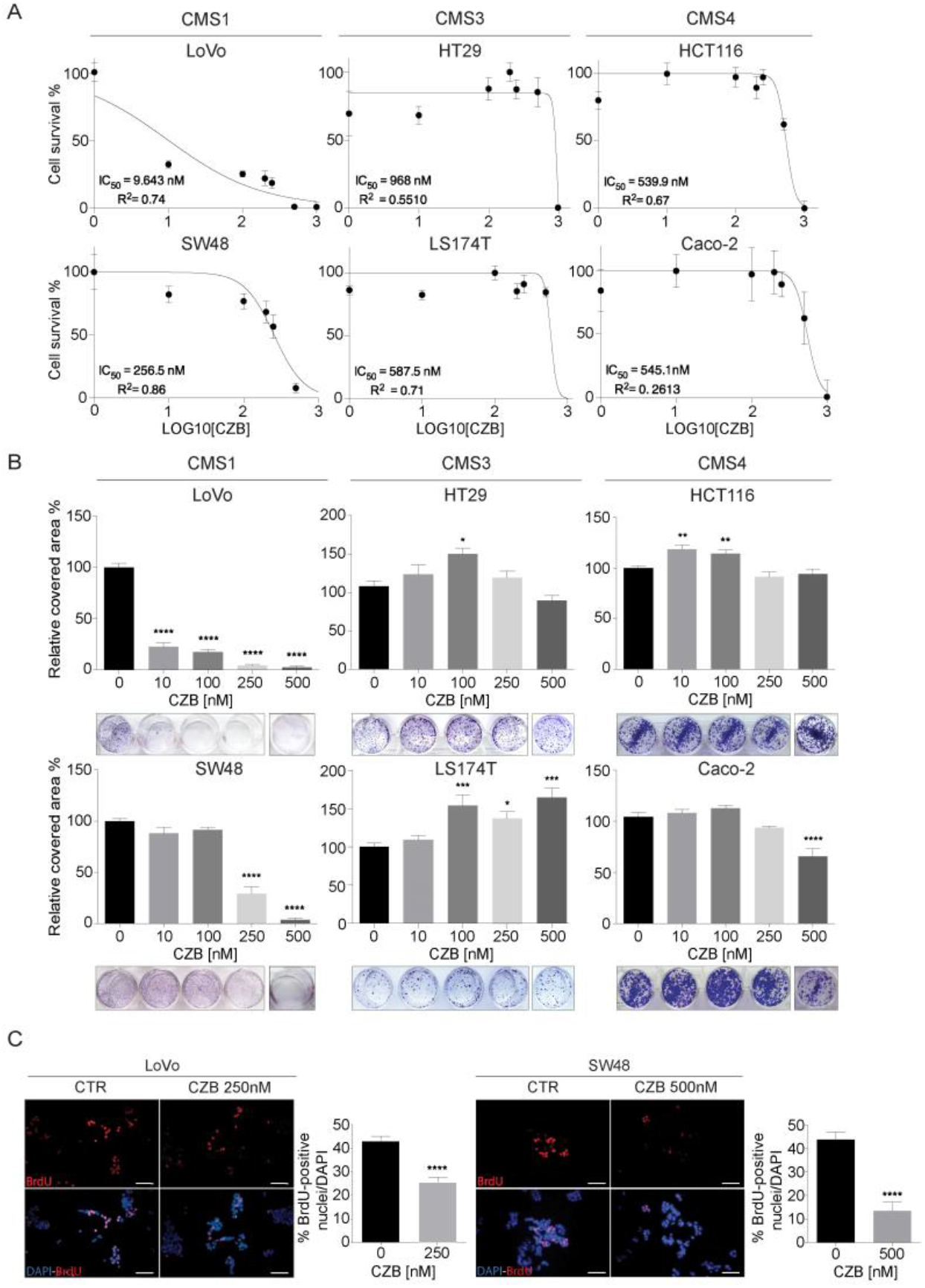
ALK INHIBITION BY CZB REDUCES PROLIFERATION OF CMS1 CELL LINES. **(A)** Cell survival assessment upon CZB addition by Alamar Blue assays. A panel of six CRC cell lines belonging to three different CMSs was tested with increasing doses of CZB. Data are reported as % of cell proliferation in relation to controls ± SD. IC50 values are shown for each cell line. CMS1 cell lines (LoVo, SW48) displayed a strong impairment in terms of cell viability after 96h of CZB exposure, detectable already at the lowest dose (CZB 10nM). CMS4 (HCT116, Caco-2) and CMS3 (HT29, LS174T) cell lines were not responsive to CZB treatment. **(B)** Colony forming assays of cell lines in (A) treated with increasing concentrations of CZB. Representative images of treated and untreated wells are shown. CZB was effective already at low doses (10nM, 250nM) only in CMS1 cells, while having no effect in CMS3 and CMS4 cells. Cell viability data are reported as % of wells covered area in relation to controls ± SD. Statistics were calculated by one-way ANOVA test. **(C)** BrdU incorporation assays of LoVo and SW48 upon CZB treatment. Representative images show BrdU-positive nuclei in red, while total DAPI-stained total nuclei are visible in blue. Decreased BrdU-positive cells were observed in LoVo and SW48 cells (CMS1) treated with CZB at the concentrations indicated. Quantifications of BrdU-positive nuclei in relation to total nuclei ± SEM is shown. Two tailed unpaired T-test was applied. Scale bar: 25μm.

**FIGURE 3.**
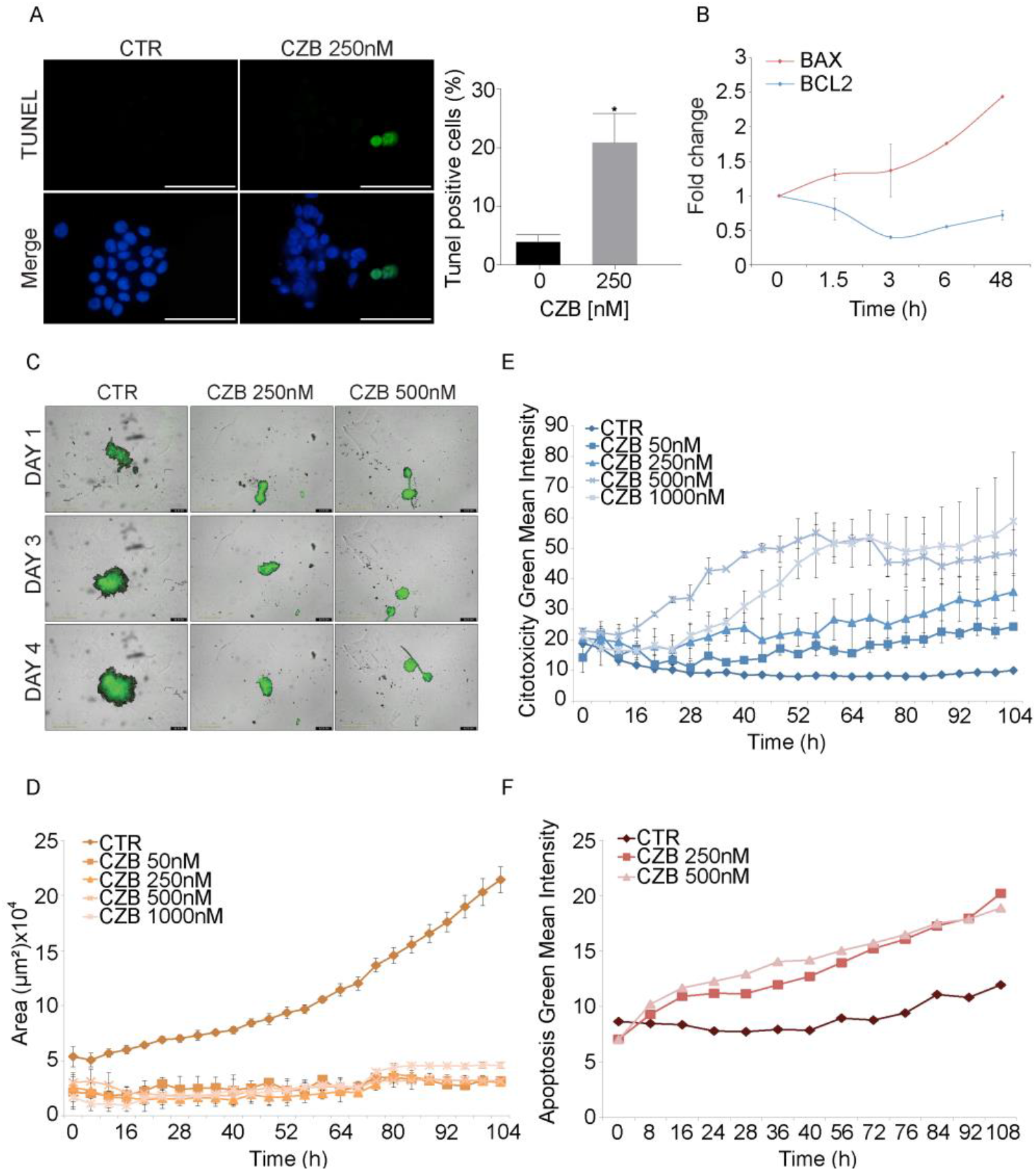
ALK INHIBITION TRIGGERS APOPTOTIC CELL DEATH IN CMS1 SUBTYPE. **(A)** Representative images (left) and quantification (right) of double-strand breaks (DSBs) assessment by TUNEL assay in LoVo cells on CZB treatment. TUNEL positive nuclei are shown in green channel, while total nuclei are visualized with DAPI in blue. ALK inhibition was shown to statistically increase DSBs levels. Two tailed unpaired T-test was applied for statistical analysis. Scale Bar: 50μm. **(B)** Determination of apoptotic markers expression by qPCR analysis performed on LoVo cells RNA, treated with CZB over a 48h time course. BCL-2 acts as an anti-apoptotic factor, while BAX as a pro-apoptotic factor. **(C)** Representative images of LoVo spheroids captured with the IncuCyte^®^ Live-Cell Analysis System. **(D, E)** Quantification of the largest brightfield object area (μm^2^) and largest brightfield object green fluorescence mean intensity (GCU) from a live-cell imaging assay performed on LoVo spheroids cultured as in (C) and treated with four different CZB doses (50nM, 250nM, 500nM, 1000nM). Spheroids were labelled with IncuCyte^®^ Cytotox Green Reagent, which showed CZB to induce cell death in a dose-dependent way also in 3D settings. Error bars refers to standard error between different replicates. **(F)** Quantification of LoVo spheroids labelled with IncuCyte^®^ Caspase-3/7 Green Reagent to detect apoptotic cells.

Time lapse imaging using a custom autofocusing method, capturing bright-field and fluorescence images every 6h for 4 days confirmed the apoptosis phenotype. Indeed, control CMS1 cells formed large aggregates within the first 3 days, with a diameter of approximately 400 μM (Fig. 3C). The IncuCyte^®^ bright-field image analysis algorithm computed the area and average size of the spheroid over time, non-invasively. ALK inhibition dramatically inhibited spheroids growth, as revealed by the impressive reduction in calculated average bright-field area (Fig. 3D). Using green fluorescent label reporters for cytotoxicity, we assessed the ALK inhibition effect on living cells. Indeed, CZB displayed a robust dose-dependent response in inducing cytotoxicity (Fig. 3E), that was strongly triggered by apoptotic pathways, as detected by Caspase-3/7 Green labelling (Fig. 3F). Time-lapse video data are available as Supp. files 1-6. In conclusion, the set of *in vitro* assays performed both in monolayer and in 3D, confirmed the enhanced antitumorigenic potential of ALK axis inhibition, specifically in CMS1.

### ALK inhibition leads to increased adhesion and cell-cell interaction

Confocal microscopy analysis of CMS1 spheroids was employed to analyze the role of ALK in the 3D cell architecture. Interestingly, on ALK inhibition, the surface organization of the spheroids displayed cell-cell tightness and increased nuclear density, with a smaller lumen cavity (Fig. 4A). Providing clear insights on the internal morphology of large spheroids or in general a thick specimen is challenging, mainly due to the poor penetration of antibodies during the staining or the hampered light penetration. Thus, to further address the evaluation of the enhanced cellular adhesion, we employed X-Clarity^TM^ Tissue Clearing System, which is able to actively extract lipids through electrophoresis to create a stable and optically transparent tissue-hydrogel hybrid that is chemically accessible for multiple rounds of antibody labeling and imaging (34).Optically cleared spheroids were evaluated by confocal microscope, that is well suited for imaging of large spheroids due to the high imaging speed and good penetration depth upon clearing (35). The microscope was equipped with silicone immersion objectives that provide long working distances and a refractive index reducing the spherical aberration caused by refractive indices mismatched between X-CLARITY mounting solution (n = 1.46) and immersion silicone oil (n = 1.40). Both ALK and e-cadherin staining were analyzed (Fig. 4B and video as Supp. file 7, 8), indicating that ALK inhibition affects the physical confinement of the nuclear geometry, by dramatically increasing cellular density on the outmost layer of the spheroids, while the cell-cell adhesion is supported by an overall enhanced e-cadherin lateral membrane localization (Fig. 4B and Supp. file 7,8). Interestingly, also the cellular localization of ALK appears regulated under CZB treatment, with increased membrane/cytosolic localization. To quantify our observations, we measured levels of vimentin, a cytoskeleton component of the intermediate filaments, by Real Time PCR. The results showed that ALK inhibition leads to a decrease in vimentin production, supporting the increased cell-cell adhesion phenotype detected by confocal microscopy (Fig. 4C).

**FIGURE 4.**
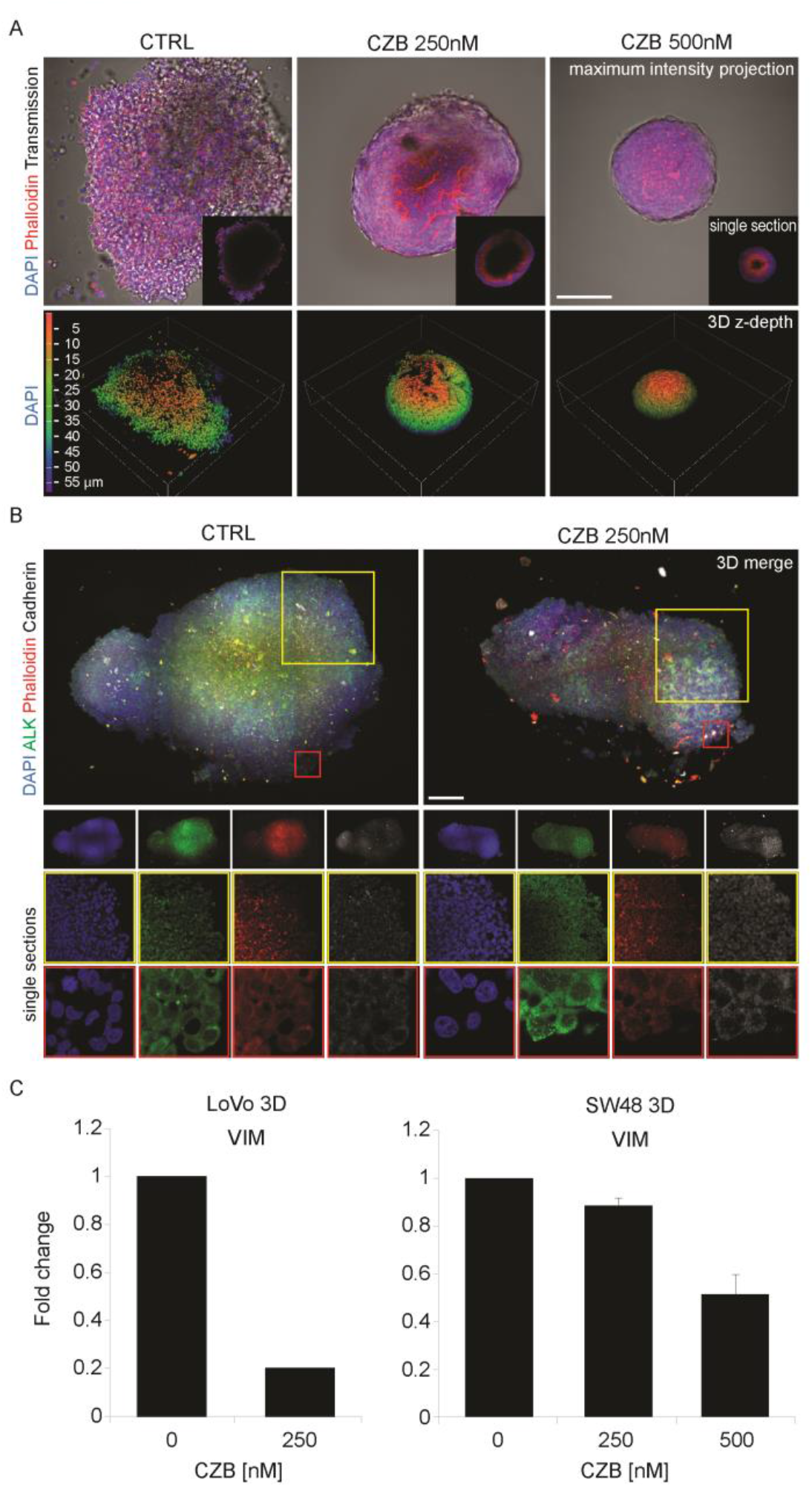
ALK INHIBITION DRIVES CELL-CELL ADHESION ENHANCEMENT IN CMS1 CELLS CULTURED IN 3D. **(A)** Confocal analysis of fixed LoVo cultured in 3D. DAPI (blue) was used to stain nuclei, while Phalloidin was used to stain actin cytoskeleton (red). The upper panel shows maximum intensity projection and single sections (insets) from control and treated spheroids. In the lower panel a volume view with 3D rendering is shown. LoVo spheroids growing in normal conditions (CTRL), exhibited an irregular morphology with detached cells. CZB was able to revert this phenotype by increasing cells organization and compactness within the sphere. Scale bar: 100μm. **(B)** Confocal analysis of clarified LoVo spheroids, stained with anti-ALK (green) and anti-E-Cadherin (grey) antibodies. Phalloidin (red) and DAPI (blue) were used to stain actin and nuclei respectively. Both 3D merges and single sections at different magnifications for each channel are shown. Scale bar: 50μm. (**C)** Determination of vimentin (VIM) expression by qPCR in LoVo and SW48 cells cultured in 3D settings and treated with CZB for 8-10 days. Data are reported in relation to untreated control.

### AKT axis is completely nullified upon ALK inhibition in CMS1

In terms of signaling output, ALK affects cell growth by activation of multiple signaling pathways, including MAPK and PI3K-AKT (36). First, we tested the ALKAL1 and ALKAL2 ligands that bind the extracellular domain of ALK, thus leading to a robust activation of the pathway (13,14). Stimulation with ALKAL1/2 ligands, alone and in combination, resulted in a strong activation of the ALK-Y1604 phosphorylation, detected already after 20 and 30 minutes of treatment (Fig. 5A), with consequent phosphorylation of the AKT axis. Notably, only a mild or no effect was recorded on the ERK axis (Fig. 5A). Next, we confirmed the specific action of CZB on ALKAL1/2 stimulating ALK activation. As expected, ALKAL2 activation of ALK and AKT was blocked on treatment with CZB (Fig. 5B). Finally, analyses of MAPK and AKT signaling in both CMS1 and CMS4 cells was performed. In line with previous findings, CZB strongly inhibited AKT activation, specifically in CMS1, while no inhibition was detected in CMS4 (Fig. 5C-D). In details, already after one hour of CZB treatment, in LoVo cells, pAKT was quenched and pERK suffered a gradual decrease, with the maximum reached after 6 hours. Surprisingly, in HCT116 cells, ALK inhibition led to a substantial increase in both pAKT and pERK levels soon after stimulation, as we detected in Fig. 5A. These data might explain the paradoxical mild positive effect we observed with CZB on the proliferation of HCT116 (Fig. 2B).

**FIGURE 5.**
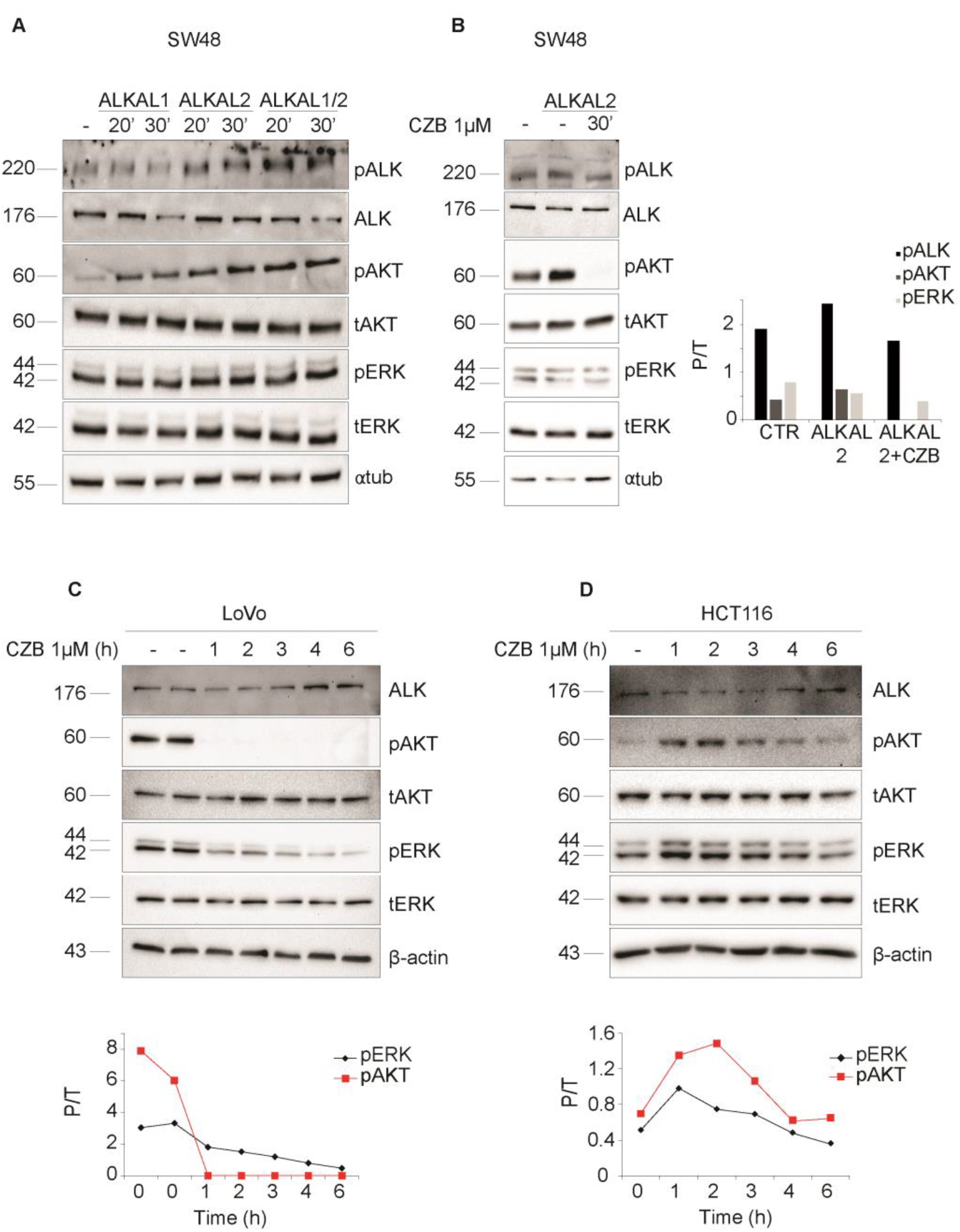
ALK DOWNSTREAM SIGNALLING PATHWAY IS ENHANCED UPON ALKALs ACTIVATION AND DAMPENED UPON CZB TREATMENT IN CMS1 CELLS. **(A)** ALK downstream signalling activity in SW48 (CMS1) cells treated with conditioned medium containing ALKAL1, ALKAL2 or their combination for either 20′ or 30′. (B) Investigation of ALK and its downstream pathway activation upon stimulation with ALKAL2 containing medium for 30′ in SW48 cells, with or without 10′ CZB pre-treatment. Quantification of pALK, pAKT and pERK is shown at right where each phosphorylated protein was normalized on the housekeeping gene and the total form, when available. **(C, D)** Western blot analysis of ALK signalling in starved LoVo (CMS1) and HCT116 (CMS4) cells, following treatment with CZB 1μM over time, from 1 to 6 hours. β-actin along with total ERK and total AKT were used as loading control. Graphs quantify pERK and pAKT over time in relation to total ERK and total AKT, respectively.

### ALK gene signature predicts survival of CMS1 patients

The inferred ability of ALK expression to predict patient survival prompted us to verify a prognostic role for the ALK-regulated gene signature. Our assumption was that the ALK signaling activation would induce a complex gene expression alteration, which could be tested in the CMSs stratified patient dataset. Because the genomic response to ALK activation is still undefined in CRC and the identification of genes that are specifically transcribed as downstream targets of this receptor remain to be addressed, we referred to a gene expression signature previously reported (GSE41635). In this data set, an unbiased transcriptome analysis was evaluated upon enforced introduction of the full-length ALK wild-type in MCF7 cells. Without filtering, approximately 1000 genes were found to be upregulated with ALK overexpression, all expressed in colon tissue. We set the proportion of false positives (PFPs) to a stringent 5%, in order to obtain a more manageable list of genes (n = 65, Supplementary Table 1). The Hochberg (step-up) multiple testing correction was applied (37) and we investigated the comprehensive correlation of this ALK signature with RFS in CRC patients. Strikingly, the ALK signature appears predictive of survival in CMS1, while a very mild (no statistically significant) association of the ALK signature for CMS4 subtype was also detected, but in this case, this subtype failed to pass the Hochberg (step-up) multiple testing correction (Fig. 6).

**FIGURE 6.**
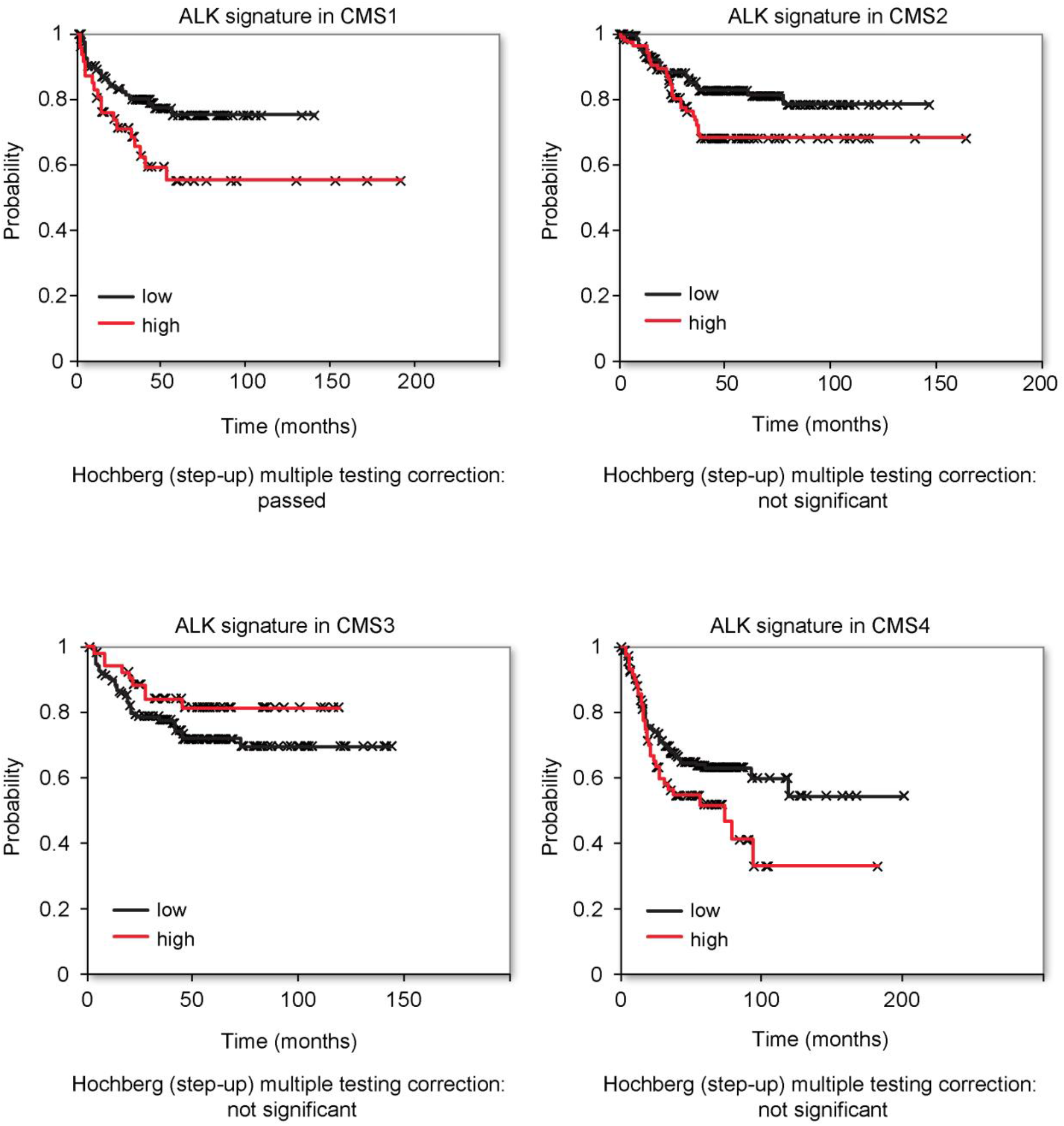
A 65-GENES ALK SIGNATURE IS PREDICTIVE OF SURVIVAL ONLY IN CMS1 COLON CANCER SUBTYPE. Bioinformatic study performed on the same dataset of patients employed for the analysis reported in Figure 1. Subjects were stratified on the basis of CMS classification and high (red line) or low (black line) expression of a signature of 65 ALK-related genes. Data were filtered according to the Hochberg (step-up) multiple testing correction. Kaplan-Meier analysis of CMS1 patients displayed a statistically significant negative correlation between RFS and ALK signature expression, in line with the results obtained for ALK. Analyses on the other CMSs were not significant.

### ALK inhibition by CZB displays therapeutic potential in patient derived organoids

We further assessed the involvement of ALK in colorectal patient-derived organoids, by testing the CZB/ALC-sensitivity in this ex-vivo model. We demonstrated that ALK inhibition induces a growth arrest only in specific patient-derived organoids, confirming the subtype specificity observed in the cancer cell lines. In details, both CZB and ALC considerably reduced CRC1430 and CRC1449 proliferation in a dose-dependent manner, but had no effect on CRC1399 organoid growth (Fig. 7A and Supplementary Fig. 2D).

**FIGURE 7.**
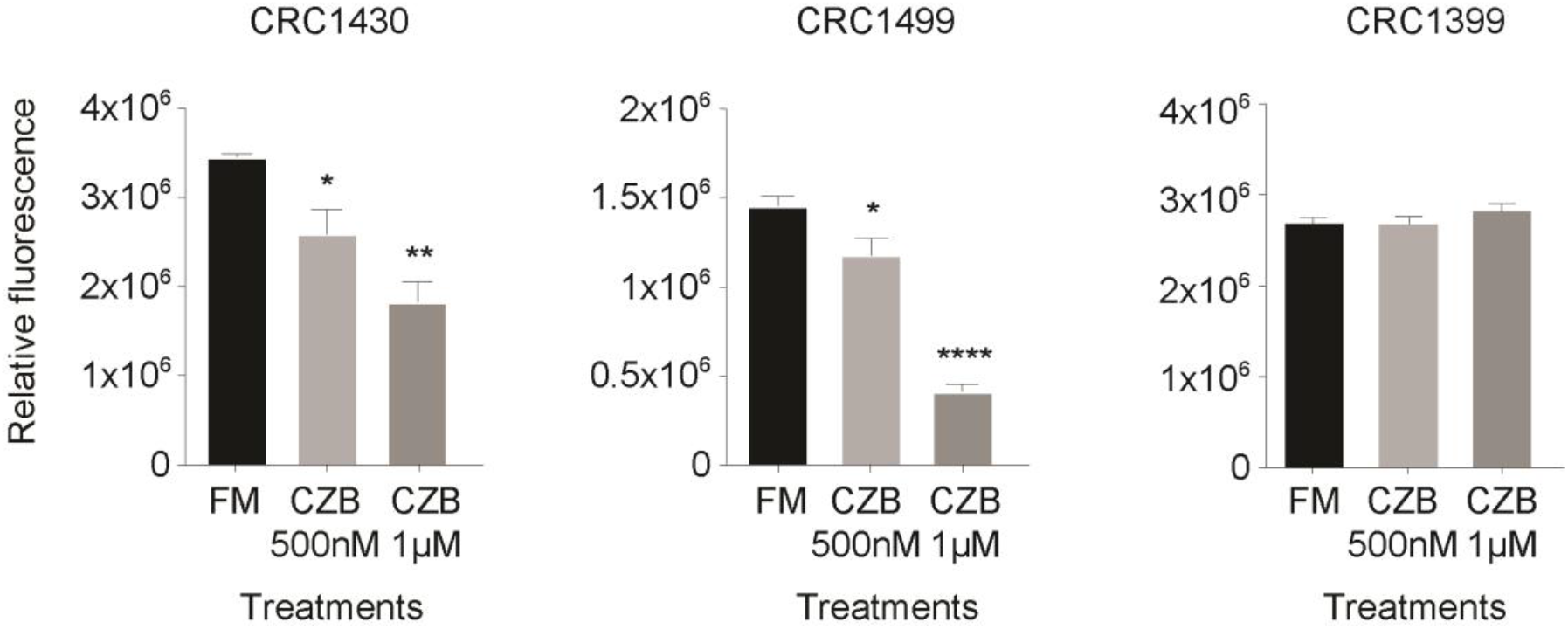
ALK INHIBITION REDUCES PROLIFERATION OF CRC PATIENT-DERIVED ORGANOIDS. Organoids proliferation assessment performed with AlamarBlue after 8/14 days of CZB treatment. Organoids were seeded in 2% BME and CZB treatment was added the following day at the indicated concentrations. Statistics were calculated by one-way ANOVA test. CZB reduced CRC1449 and CRC1430 growth, while having no effect on CRC1399.

## DISCUSSION AND CONCLUSION

The CMS classification has proven to be a useful prognostic factor for early stage colorectal cancer (6). In this study, we assessed the impact of ALK inhibition on CMS outcomes. This hypothesis-generating study stemmed from the finding that ALK receptor mRNA abundance, as well as ALK pathway activation (as measured with a 65 gene ALK signature), are associated with survival in CMS1 patients subtype while displaying no effect in the other CMS subtypes. Notably, evidence for an involvement of ALK in colorectal cancer has already been suggested, mostly arguing for a cross-talk with EGFR and revealing the presence of tumours harbouring both tyrosine kinase receptors alteration or ALK gene amplification as a poor prognostic factor (23,38). Our data on the *in vitro* sensitivity to ALK inhibition were congruent in multiple cell lines and settings, from proliferation to soft agar and three-dimensional assays, which are known to be closer to tumours *in vivo*. The cellular system we applied reliably recapitulates the CMS1 subtype and strongly supports the therapeutic inhibition of ALK in this specific CRC subset. Interestingly, four additional cell lines belonging to CMS 3 and 4 subtypes, confirmed a complete lack of response. The employment of *in vitro* models led to the detection of a signalling involving AKT axis downstream of ALK activation. Finally, a functional role for ALK inhibition in CMS1 was confirmed by a comprehensive imaging analysis of the 3D tumour formation, with state of the art clearing technology and silicone immersion objectives that provide superior depth, high resolution performance and minimize spherical aberration. From a clinical point of view, metastatic CRC maintains a unique dependency on the receptors of the ErbB family, which are the target of few monoclonal antibodies that robustly downregulate EGFR signaling. The results presented here suggest a CMS1 dependence on ALK pathway for growth and survival. We are aware that experiments with cell lines cannot recapitulate the wide complexity of tumors described on a population basis. Nevertheless, the cell line work was tightly supported and in line with the patients derived data and Kaplan-Meier profiles. The observation that *ALK* is relevant in a small percentage of CRC patient is in agreement with previous reports generated on unselected colorectal cancers specimens (22,23). The unexpected novel finding here is the association of ALK with the CMS1 and consequent therapeutic implications for this patient group. Previously ALK translocation has been associated to high MSI CRC, which in our analyses cover the CMS1 subtype (39). Interestingly, these data find support in some clinical activity registered for the entrectinib, in patients with *ALK* translocation in gastrointestinal cancer, who had never been treated with agents designed to target those alterations. Definitely, this work highlights the growing assumption that integration of clinical data with functional analyses of preclinical models can lead to rational therapies aimed at delaying or halting the occurrence of CRC in specific CMSs. To sum up, here we suggest that ALK pathway activation may represent a possible target for personalized therapy, in CRC patients who currently have limited treatment options.

## MATERIALS AND METHODS

### CELL CULTURES

Experiments were performed using several colorectal cancer (CRC) cell lines and patient-derived organoids (n=3), kindly provided by Prof. Livio Trusolino laboratory. LoVo, HCT116, Caco-2, HT29 and LS174T were grown in Dulbecco’s Minimal Essential Medium (DMEM) high-glucose, while SW48 were grown in Leibovitz medium (L-15). Media were supplemented with 10% fetal bovine serum (FBS) and 1% penicillin-streptomycin. Organoids were cultured in 100% Cultrex^®^ RGF BME (R&D Systems, Minneapolis, USA) covered with DMEM/F12 (1% penicillin-streptomycin and 2mM L-Glutamine) supplemented with B-27 1X, N-2 1X, 1mM N-acetylcysteine and EGF 10ng/ml. Cells and organoids were cultured in a humidified 37°C incubator with 5% CO2, except SW48 cell line, which has been kept in dioxide-free conditions as required for L-15 medium maintenance. All cell lines were routinely tested for *Mycoplasma* contamination by PCR. Caco-2, LoVo and HCT116 were validated with the Promega, PowerPlex 21 PCR Kit, from the external service Eurofins Medigenomix, Ebersberg.

### TRANSFECTION

HEK293 cells were grown in the absence of antibiotics and, at about 90% confluence, were transfected using Lipofectamine 2000, according to manufacturer’s instructions (Invitrogen, Carlsbad, USA). ALKAL1/2 expressing plasmids have been previously described (13). Both plasmids and Lipofectamine were diluted in Opti-MEM before being mixed together and, after 20 minutes of incubation at room temperature, added to the cells. After 12 hrs medium was replaced with serum-free DMEM and 48 hrs after transfection medium was harvested, centrifuged and stored at −80°C for subsequent assays.

### PROLIFERATION ASSAY

For AlamarBlue (resazurin) proliferation assays, cells were seeded in full medium in 96- well plates. Quantification of initial time (time 0) was performed the following day, adding 200μM of AlamarBlue in each well, as previously reported (31). Cells were then treated according to the experiment and proliferation was measured again after 96 hours following the same procedure previously described. Data were analysed as previously reported (30).

### COLONY FORMING ASSAY

2.000 (Caco-2, HCT116, HT-29, LS174T), 5000 (LoVo) or 10000 (SW48) cells were seeded in 12-well plates and treated the following day according to the experiment. After ten days cells were fixed and stained as previously described (30). For each well, several pictures were taken and percentage of the area covered by cells was quantified using ImageJ software.

### BrdU INCORPORATION ASSAY

Cells were seeded in full medium on coverslips in a 12-well plate and, after 2 days, treated for 24/48h according to the experiment. At the end of the treatment, cells were incubated with 1 μg/ml BrdU for 20 or 30′. Cells were then firstly fixed in 4% PFA and later incubated in 2N HCl at 37°C for 15′, causing DNA hydrolysis. After treating with 0.2% Triton and blocking with 1% bovine serum albumin (BSA, Sigma-Aldrich, St. Louis, USA), 3ug/ml anti-BrdU primary antibody was added O/N. The next day, cells were incubated with 1:100 Cy3 secondary antibody and 1:1000 DAPI and the coverslip was finally mounted onto the slide. Images were taken using the Olympus BH-2 CCD microscope. Positive cells were counted using the ImageJ software.

### TUNEL ASSAY

Cells were seeded in full medium on coverslips in a 12-well plate and, after 1 day, treated for 48h according to the experiment. After fixation in 4% PFA, cells were permeabilised with 0.2% Triton and treated for 1h with the TUNEL reaction mixture (enzyme and label solution), except for the negative control where the enzyme was not added. Lastly, cells were incubated with 1:1000 DAPI and the coverslip was mounted onto the slide. Images were taken using the Olympus BH-2 CCD microscope, detecting fluorescence in the range of green. Positive cells were counted using the ImageJ software.

### CONFOCAL MICROSCOPY AND SPHEROIDS CLEARING

Confocal microscopy was used to investigate the inner structure of spheroids and potential phenotypical changes in response to drugs treatment. Spheroids from 3D culture obtained as reported in Supplementary Methods section were collected, fixed and stained according to the procedure previously described (30). See the list of reagents in Supplementary Information for additional details about fluorescent dyes. When necessary, spheroids clearing was performed prior to staining and microscope observation. For this purpose, samples were incubated in the X-CLARITY Hydrogel-Initiator solution (Logos Biosystems, Inc. Anyang-Si, Gyunggi-Do, South Korea) according to manufacturer’s instructions. Samples were polymerized with the X-Clarity^TM^ Polymerization System (Logos Biosystems) for 3 hrs with a vacuum of 90 kPa and a temperature of 37°C. After polymerization, spheroids were gently shaken for 1′, rinsed several times with PBS and stored at 4°C. Subsequently, samples-hydrogel hybrid were immersed in an Electrophoretic Tissue Clearing Solution (Logos Biosystems) and placed for 8 hrs in the X-Clarity^TM^ Tissue Clearing System (Logos Biosystems) with the following settings: current 0.8 A, temperature 37°C, pump speed 30 rpm. After several washes in PBS, spheroids were then incubated with 4% BSA (Sigma-Aldrich) to avoid unspecific bindings prior to incubation with primary and secondary antibodies. The complete list of the used antibodies can be found in the Supplementary Information section. Samples were then mounted with a mixture of X-CLARITY mounting solution (Logos Biosystems) and 1,4-Diazabicyclooctane (DABCO) (Sigma-Aldrich). Confocal imaging from non-clarified samples was performed with a Nikon A1-R confocal laser scanning microscope, equipped with a 20× 0.7 NA objective and with 405 and 561 nm laser lines to excite DAPI and red fluorescence signals. Fluorescent images from clarified spheroids were visualized and imaged under FV300 fluorescent confocal microscope (Olympus, Shinjuku, Japan) equipped with a 30x (silicon immersion, 1.05 NA) and 60× (silicon immersion, 1.30 NA) super apochromat objectives and with 405, 488, 561 and 647 nm laser lines. Z-stacks were collected at optical resolution of 210 nm/pixel, stored at 12-bit with 4096 different gray levels, pinhole diameter set to 1 Airy unit and z-step size to 1μm. The data acquisition parameters were setup in fixed manner, such as laser power, gain in amplifier and offset level. Confocal images were processed using Richardson-Lucy *deconvolution* algorithm. Volume view with 3D rendering was carried out using the NIS Elements Advanced Research software (Nikon, Shinagawa, Jaoan).

### INCUCYTE^®^3D SINGLE TUMOR SPHEROID ASSAYS

Spheroids growth was followed in time-lapse by means of IncuCyte S3^®^ Live-Cell Analysis system (Essen Bioscience, Ann Arbor, USA). For live imaging assays, cells were seeded in 96-well plates over a layer of 0.6% agar as performed for spheroid assays. Fluorescent reagents for detection of cytotoxicity and apoptosis, designed specifically for IncuCyte S3^®^ experiments by Essen Bioscience, were added at the time of seeding, according to manufacturer’s instructions. More details about the dyes for live-cell experiments can be found in the Fluorescent Probes List in Supplementary Information section. Analyses were performed using IncuCyte S3^^®^^ Software (v2018C).

### WESTERN BLOT

Starved and treated cells were lysed using RIPA buffer supplemented with a protease inhibitor cocktail (P8340, Sigma-Aldrich, 1:100) and 1mM Na_3_VO_4_. Protein concentration in the supernatants was determined by DC Protein Assay (Bio-Rad, Hercules, USA), using bovine serum albumin as the standard. Protein samples were separated by SDS-PAGE and then transferred to nitrocellulose membranes (Amersham™ Protran™ Premium 0.45μm 300mm × 4m, GE Healthcare, USA). Membranes were blocked for 1 hour using 3% BSA (Sigma-Aldrich) in TBS-T (0.1% Tween-20) and incubated O/N (4°C) with primary antibodies diluted in the blocking solution. Finally, membranes were incubated with horseradish peroxidase-coniugated secondary antibodies against mouse or rabbit for 1 hour and protein presence was detected by chemiluminescent reaction (Amersham^TM^ ECL^TM^ Detection Reagents). Bands quantification was performed using Image Lab software. Details about the antibodies employed for proteins detection are reported in Supplementary Information.

### qRT-PCR

Total RNA was extracted from starved cells using QIAzol^^®^^Lysis Reagent (QIAGEN, Venlo, Netherlands) according to manufacturer’s instructions. qRT-PCR analyses were then performed on cDNA using the iTaq universal SYBR Green Supermix (Bio-Rad). A complete list of the primers employed is reported in the Supplementary Information section. Gene expression data from qRT-PCRs were determined as ΔCq (Quantification Cycle) normalized on the expression of β_2_-microglobulin housekeeping gene.

### STATISTICS

The statistical analyses were performed by using Prism version 6 (GraphPad Software, Inc). Both T-test and one-way ANOVA were used to test significance of the assays. The details of the applied statistical test are reported in the figure legends. * Pvalue < 0.05; ** Pvalue < 0.01; *** Pvalue < 0.001; **** Pvalue < 0.0001

### Author contributions

ML, RP and GD were involved in designing research studies. MM, MS, VG, DR, SS, CC and MF and were responsible in conducting experiments, acquiring data and analyzing data. BG performed the bioinformatic data analyses. MM, MS and ML wrote the manuscript. RP provided reagents and revised the manuscript.

## Acknowledgments

We thank Dr. Luca Cevenini (Olympus Italia) and Dr. Mario Barlocco (Twin Helix) for technical assistant. This research was funded by PRIN2017, grant number 2017TATYMP_002, Fondazione Carisbo, Fondazione Cariplo and AlmaIdea2018.

## Notes

### Competing Interest Statement

The authors have declared no competing interest.

