## Supplemental Data for "Pharmacological inhibition of the ALK axis elicits therapeutic potential in Consensus Molecular Subtype 1 colon cancer patients"

**LIST OF REAGENTS**

ANTIBODIES LIST

| ANTIBODY | SOURCE | CATALOG CODE |
|---|---|---|
| Phospho-AKT (Ser473) (D9E) XP Rabbit monoclonal antibody | Cell Signalling Technology | #4060 |
| AKT Rabbit polyclonal antibody | Cell Signalling Technology | #9272 |
| MAP Kinase activated (diphosphorylated ERK 1/2) Mouse monoclonal antibody | Sigma-Aldrich | M8159 |
| ERK 2 (D-2) Mouse monoclonal antibody | Santa Cruz Biotechnology | sc-1647 |
| ALK (F-12) Mouse monoclonal antibody | Santa Cruz Biotechnology | sc-1647 |
| pALK (Tyr1604) Rabbit monoclonal antibody | Cell signalling Technologies | #3341 |
| β-Actin (C4) Mouse monoclonal antibody | Santa Cruz Biotechnology | sc-47778 |
| α-Tubulin (B-7) Mouse monoclonal antibody | Santa Cruz Biotechnology | sc-5286 |
| E-cadherin (G-10) Mouse monoclonal antibody | Santa Cruz Biotechnology | sc-8426 |
| Anti-Mouse/Anti-Rabbit EnVision+ System- HRP Labelled Polymer | Dako | K4001/K4003 |
| Anti-BrdU | DSHB | #G3G4 (AB_2618097) |

FLUORESCENT PROBES LIST

| FLUORESCENT PROBE | SOURCE | CATALOG CODE |
|---|---|---|
| Alexa Fluor™ Phalloidin | Thermo Fisher Scientific | #A12379 |
| DAPI | Sigma-Aldrich | #D9542 |
| IncuCyte® Caspase-3/7 Green Apoptosis Assay Reagent | Essen BioScience Inc | Cat. No. 4440 |
| IncuCyte® Cytotox Green Reagent for Counting Dead Cells | Essen BioScience Inc | Cat. No. 4633 |

PRIMERS FOR qRT-PCR LIST

| TARGET TRANSCRIPT | SEQUENCE | SOURCE |
|---|---|---|
| BCL2 | FW: ATGTGTGTGGAGAGCGTCAACC | Sigma-Aldrich |
| | RW: TGAGCAGAGTCTTCAGAGACAGC | Sigma-Aldrich |

| | | |
|---|---|---|
| BAX | FW: GGGACGAACTGGACAGTAACA | Sigma-Aldrich |
| | RW: CCGCCACAAAGATGGTCA | Sigma-Aldrich |
| VIM | FW: GGAAACTAATCTGGATTCACTC | Sigma-Aldrich |
| | RW: CATCTCTAGTTTCAACCGTC | Sigma-Aldrich |
| B2M | FW: TGCCTGCCGTGTGAACCATGT | Sigma-Aldrich |
| | RW: TGCGGCATCTTCAAACCTCCATGA | Sigma-Aldrich |

**SUPPLEMENTARY METHODS**

### SPHEROID ASSAY

Spheroids formation was prompted by growing cells in low-attachment conditions. For this purpose, wells from 6-well plates were coated as previously described (29) Cells were then seeded in full medium supplemented with proper treatments and with EGF 10ng/mL, to promote spheroids formation, when necessary. Spheroids volume was assessed as previously reported by means of ImageJ Software (29).

### SOFT AGAR ASSAY

Cells were grown embedded into a layer of sterile agar 0.3% in full medium plus proper treatments, on wells previously coated with agar 0.6% as previously described for spheroid assays. A thin growth medium layer was added weekly on top of agar layer to prevent desiccation. After 15 days-1 month, depending on each cell line, photos of 3D colonies were taken using inverted microscope (Leitz Labovert FS, code 515). Embedded cells were then incubated with PFA4% for fixation and stained with Giemsa 3-5% in PBS 1X. Photographs of each well were taken and colonies' number was assessed with ImageJ Cell Counter plugin.

### FLUORESCENT IN SITU HYBRIDIZATION

Samples consisted of small formalin-fixed and paraffin-embedded cell pellets. FISH assay was performed using the Vysis ALK Break Apart FISH Probe Kit (Abbott). This break-apart FISH test is based on a mixture of two probes hybridizing to the proximal (3', orange-labeled probe) and distal (5', green-labeled probe) to the ALK breakpoint cluster region. At least 50 non-overlapping nuclei were scored for each specimen by a trained technologist and a pathologist. Cells positive for rearrangement are defined by two main patterns: I) a "split pattern", with 3' and 5' break apart signals at a distance of two times the diameter of the largest signal; II) a "5' deletion pattern", showing one fusion signal and an isolated 3' orange signal (without the corresponding 5' green signal). A case was considered FISH positive for ALK rearrangements when at least 15% of tumour cells showed any split or any 5’ deletion pattern.

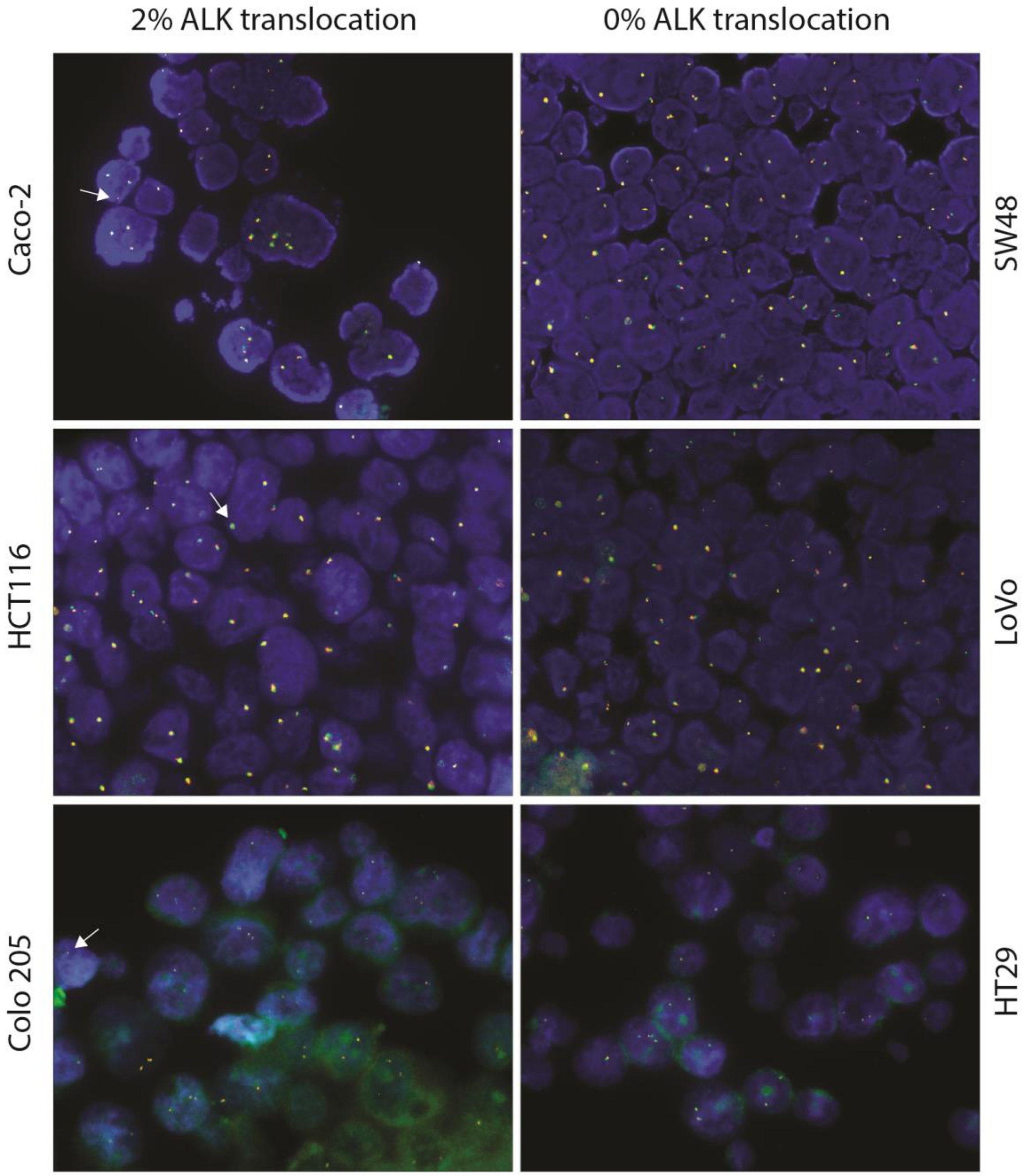


**Supplementary Figure 1 - ALK STATUS IN A PANEL OF CRC CELL LINES BELONGING TO DIFFERENT CONSENSUS MOLECULAR SUBTYPES**

Break-apart FISH assay performed on cell pellets fixed with formalin and embedded in paraffin. The lack of significant *ALK* aberrant rearrangements was assessed in all the cell lines tested.

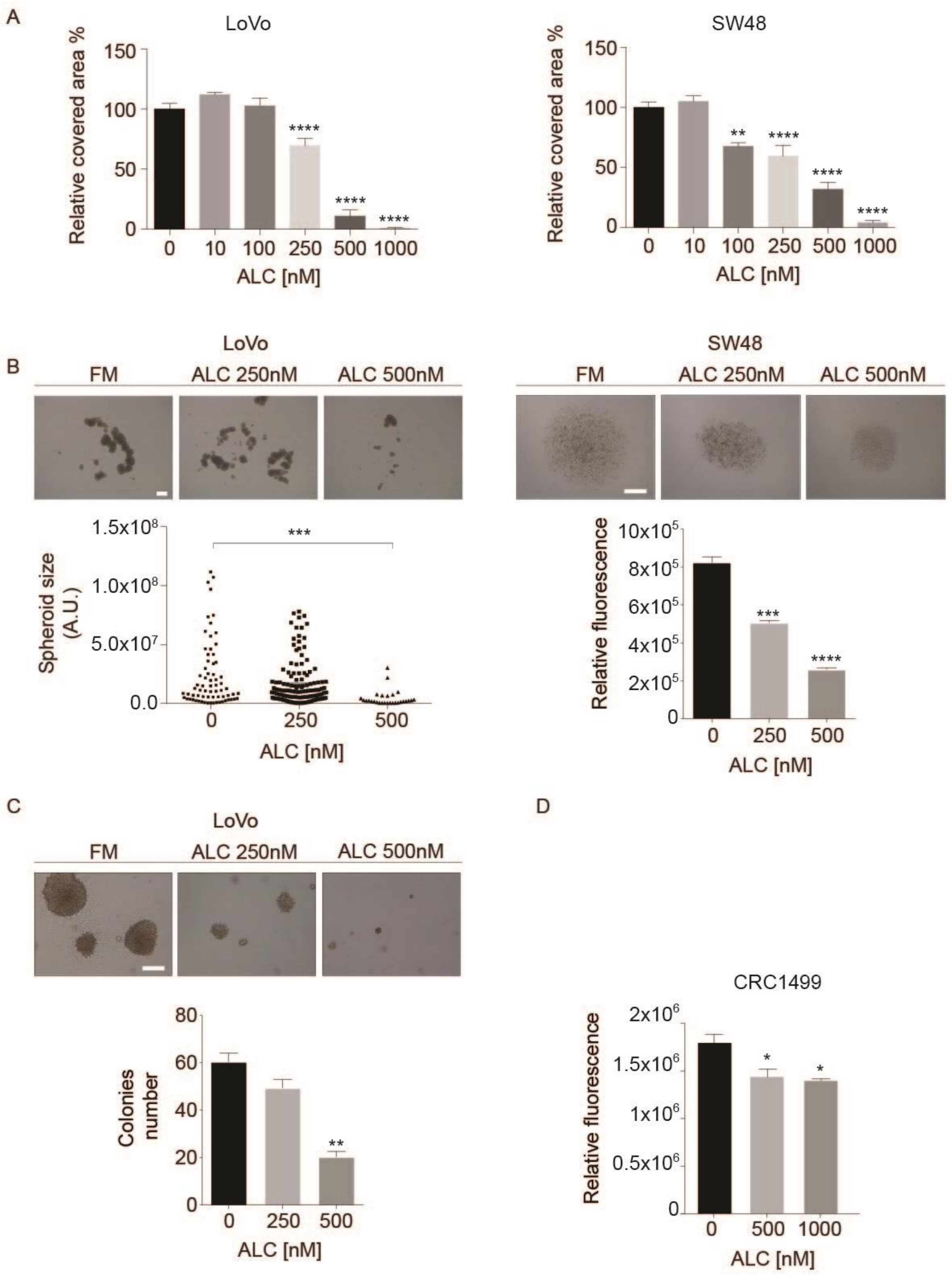


**Supplementary Figure 2 - ALK INHIBITOR ALECTINIB REDUCES PROLIFERATION OF CRC PATIENT-DERIVED ORGANOIDS AND CMS1 CELLS IN 2D AND 3D SETTINGS**

**(A)** Colony forming assays of LoVo and SW48 cells treated with increasing ALC concentrations. Statistic was performed by one-way ANOVA. In both cell lines, ALC is effective already at 250nM. **(B)** Spheroid assay of LoVo and SW48 cells treated with ALC. Spheroids formation was not observed in SW48, which tend to grow in dispersed aggregates. For LoVo cells, spheroid's size was calculated through the formula (minor axis x major axis$_{2}$) x 2 and related to controls, while SW48 cells' growth in 3D was measured using Alamar Blue assay. Statistic was calculated by one-way ANOVA. ALC is clearly able to dampen SW48 and LoVo cells when cultured in lack of adhesion. Scale bars: SW48: 2000 µm. LoVo: 750 µm. **(C**) Soft-agar assay of LoVo cells treated with ALC. Representative images of 3D colonies are reported. Colonies were counted after Giemsa staining by means fo ImageJ software. Statistic was calculated by one-way ANOVA. ALK inhibition by ALC impairs proliferation of LoVo 3D colonies growing embedded in 0.3% agar. Scale bar: 250 µm. **(D)** CRC1449 proliferation assessment performed with AlamarBlue after 11 days of ALC treatment. Patient-derived organoid was seeded in FM + 2% BME and treatments were added the following day. Statistic was calculated by one-way ANOVA test. ALC was able to reduce CRC1449 growth over time.

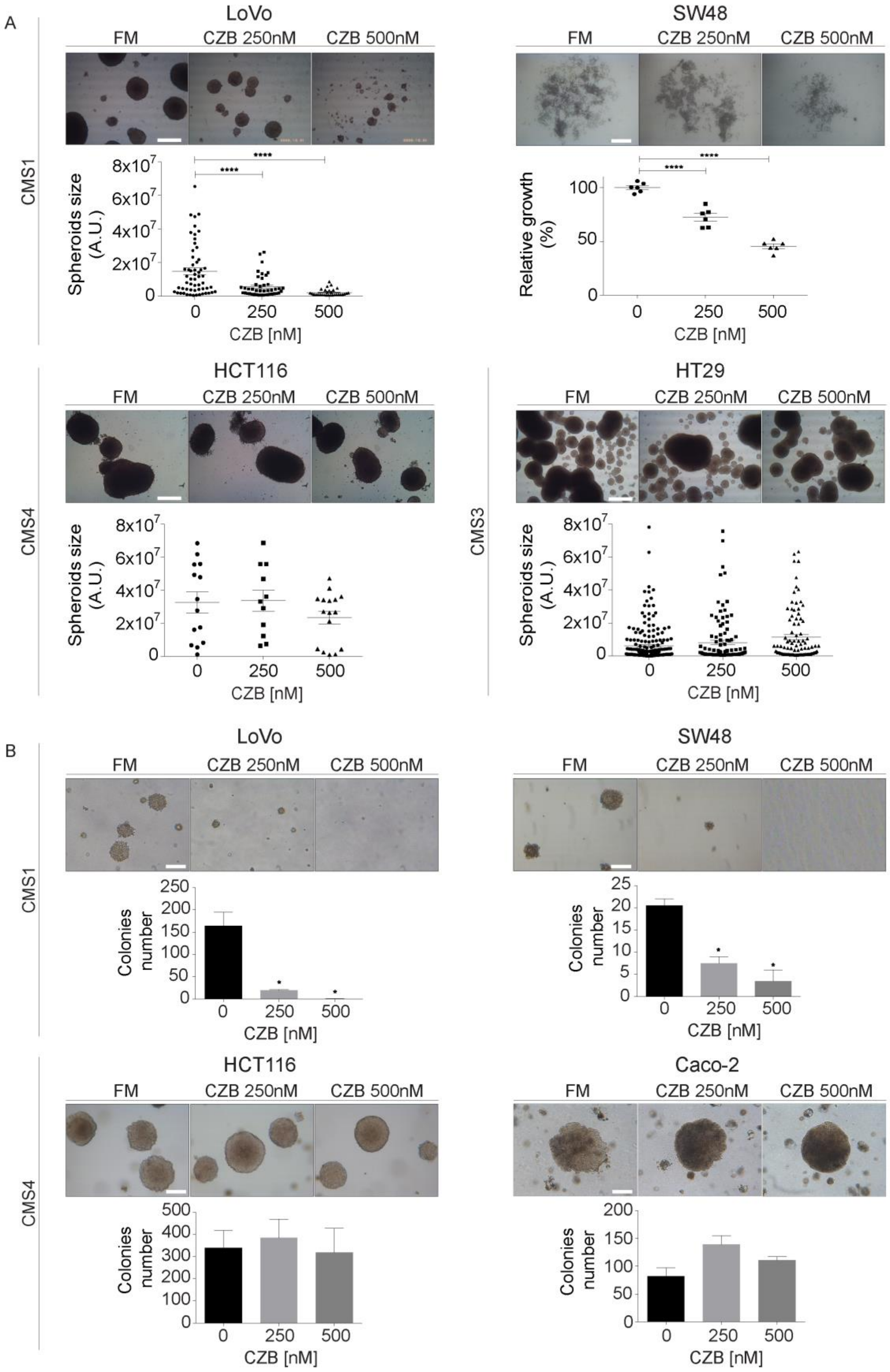


**Supplementary Figure 3 - ALK INHIBITION SIGNIFICANTLY IMPACTS ON CMS1 SPHEROIDS GROWTH**

**(A)** Spheroid-forming assays of four cell lines representative of different subtypes. HCT116, LoVo and HT29 spheroids' size was calculated through the formula (minor axis x major axis$_2$) x 2 and related to controls. In case of SW48 cells, proliferation was measured through Alamar Blue assay. Statistic was calculated by one-way ANOVA. Scale bars: HCT116, LoVo, HT29: 250 µm. SW48: 2000 µm. **(B)** Soft-agar assays of four cell lines belonging to different CMSs. For each cell line, representative images are reported to show the phenotype and size of colonies. A semi-quantitative value of cells invasiveness was derived by counting the number of colonies after Giemsa staining, as showed in column charts. Statistic was calculated by one-way ANOVA. Scale bar: 250 µm.
