## Supplementary Table 1 for "Pharmacological inhibition of the ALK axis elicits therapeutic potential in Consensus Molecular Subtype 1 colon cancer patients"

| probe set | gene | gene.index | RP/Rsum | FC:(class1/class2) | pfp | P.value |
|---|---|---|---|---|---|---|
| 209987_s_at | ASCL1 | 19110 | 1,732 | 1,21E+22 | 9,64E-05 | 1,79E-09 |
| 209988_s_at | ASCL1 | 19111 | 2 | 1,91E+19 | 7,32E-05 | 2,71E-09 |
| 208211_s_at | ALK | 17388 | 2,449 | 5,61E+19 | 8,67E-05 | 4,82E-09 |
| 201215_at | PLS3 | 10509 | 4,899 | 1,99E+15 | 0,0004078 | 3,02E-08 |
| 226618_at | FLJ25076 | 35373 | 5,831 | 3,61E+09 | 0,0005055 | 4,68E-08 |
| 228046_at | LOC152485 | 36784 | 8,775 | 1,67E+10 | 0,001147 | 1,28E-07 |
| 228640_at | PCDH7 | 37376 | 9,487 | 1,07E+09 | 0,001187 | 1,54E-07 |
| 226764_at | | 35518 | 9,798 | 92235803 | 0,001122 | 1,66E-07 |
| 228915_at | DACH1 | 37647 | 10,68 | 447127522 | 0,001224 | 2,04E-07 |
| 227276_at | PLXDC2 | 36025 | 11,53 | 82774305 | 0,001324 | 2,45E-07 |
| 220615_s_at | MLSTD1 | 29469 | 13,75 | 2,80E+07 | 0,001822 | 3,71E-07 |
| 209387_s_at | TM4SF1 | 18527 | 14,7 | 1860691 | 0,001953 | 4,34E-07 |
| 213110_s_at | COL4A5 | 22093 | 17 | 3425104 | 0,002532 | 6,10E-07 |
| 204915_s_at | SOX11 | 14163 | 18,49 | 1805857 | 0,002859 | 7,42E-07 |
| 223918_at | ACSL6 | 32723 | 20,2 | 932487 | 0,003273 | 9,10E-07 |
| 204971_at | CSTA | 14218 | 22 | 394484 | 0,003735 | 1,11E-06 |
| 228635_at | PCDH10 | 37371 | 22,98 | 112454 | 0,003886 | 1,22E-06 |
| 209735_at | ABCG2 | 18863 | 25,92 | 137123 | 0,004838 | 1,61E-06 |
| 236297_at | --- | 44939 | 27,66 | 52517 | 0,005314 | 1,87E-06 |
| 219440_at | RAI2 | 28305 | 27,93 | 242179 | 0,005161 | 1,91E-06 |
| 239195_at | --- | 47797 | 32,31 | 57086 | 0,006848 | 2,67E-06 |
| 205696_s_at | GFRA1 | 14940 | 32,53 | 43409 | 0,006637 | 2,71E-06 |
| 226612_at | FLJ25076 | 35368 | 32,62 | 37319 | 0,006389 | 2,72E-06 |
| 238575_at | OSBPL6 | 47183 | 33,05 | 19972 | 0,006306 | 2,80E-06 |
| 202478_at | TRIB2 | 11755 | 38,34 | 16871 | 0,008474 | 3,93E-06 |
| 206440_at | LIN7A | 15671 | 39,33 | 13533 | 0,008631 | 4,16E-06 |
| 203413_at | NELL2 | 12683 | 39,75 | 12420 | 0,008511 | 4,26E-06 |
| 1553995_a_at | NT5E | 1231 | 51,23 | 4043 | 0,01452 | 7,53E-06 |
| 204584_at | L1CAM | 13836 | 51,93 | 4491 | 0,01445 | 7,76E-06 |
| 218793_s_at | SCML1 | 27661 | 52,44 | 3268 | 0,01428 | 7,94E-06 |
| 225664_at | COL12A1 | 34431 | 59,4 | 3002 | 0,01825 | 1,05E-05 |
| 204724_s_at | COL9A3 | 13974 | 62,35 | 2491 | 0,0197 | 1,17E-05 |
| 217284_x_at | SERHL | 26180 | 63,71 | 2654 | 0,02004 | 1,23E-05 |
| 233998_x_at | --- | 42664 | 64,5 | 257681 | 0,01999 | 1,26E-05 |
| 209985_s_at | ASCL1 | 19108 | 64,81 | 2581 | 0,01963 | 1,27E-05 |
| 238455_at | --- | 47066 | 65,35 | 884,1 | 0,01944 | 1,30E-05 |
| 223605_at | | 32417 | 67,28 | 2293 | 0,02018 | 1,38E-05 |
| 235913_at | LOC400713 | 44560 | 70,99 | 183,8 | 0,02215 | 1,56E-05 |
| 212154_at | SDC2 | 21152 | 73,83 | 1457 | 0,02354 | 1,70E-05 |
| 217276_x_at | SERHL2 | 26172 | 74,3 | 1688 | 0,02328 | 1,73E-05 |
| 213768_s_at | ASCL1 | 22741 | 74,99 | 1939 | 0,02318 | 1,76E-05 |
| 209083_at | CORO1A | 18229 | 75,58 | 1338 | 0,02302 | 1,79E-05 |
| 230163_at | LOC143381 | 38882 | 76 | 1364 | 0,02277 | 1,81E-05 |
| 203423_at | RBP1 | 12693 | 76,75 | 494,3 | 0,02274 | 1,85E-05 |
| 219737_s_at | PCDH9 | 28600 | 80,83 | 1331 | 0,02494 | 2,08E-05 |
| 225667_s_at | --- | 34434 | 80,87 | 1034 | 0,02443 | 2,08E-05 |
| 225651_at | UBE2E2 | 34418 | 81,24 | 1113 | 0,02415 | 2,10E-05 |
| 1557533_at | --- | 3655 | 81,85 | 1161 | 0,02404 | 2,14E-05 |
| 217146_at | JRK | 26045 | 87,43 | 649,8 | 0,02725 | 2,47E-05 |
| 232315_at | LOC400713 | 41002 | 88,25 | 609 | 0,02727 | 2,53E-05 |
| 232210_at | --- | 40899 | 88,81 | 934,5 | 0,02711 | 2,56E-05 |

| | | | | | | |
|---|---|---|---|---|---|---|
| 214540_at | HIST1H2BO | 23503 | 89,22 | 231,4 | 0,02686 | 2,59E-05 |
| 223883_s_at | STK31 | 32692 | 89,39 | 942,2 | 0,02646 | 2,60E-05 |
| 216456_at | | 25377 | 92,61 | 823,1 | 0,02808 | 2,81E-05 |
| 207826_s_at | ID3 | 17021 | 94,23 | 734,7 | 0,02866 | 2,92E-05 |
| 218312_s_at | ZSCAN18 | 27187 | 95,07 | 889,9 | 0,0287 | 2,98E-05 |
| 214807_at | | 23763 | 99,14 | 566,4 | 0,03093 | 3,27E-05 |
| 206504_at | CYP24A1 | 15734 | 105,5 | 537,7 | 0,03488 | 3,75E-05 |
| 222747_s_at | SCML1 | 31570 | 112,8 | 551,1 | 0,03974 | 4,34E-05 |
| 234715_at | GOLGA2LY1 | 43373 | 115,1 | 249,4 | 0,04081 | 4,54E-05 |
| 242660_at | C10orf112 | 51228 | 118,1 | 431,1 | 0,04247 | 4,80E-05 |
| 1561501_s_at | LOC348180 | 5972 | 121,4 | 356,2 | 0,04444 | 5,11E-05 |
| 204638_at | ACP5 | 13890 | 125,1 | 368,4 | 0,04673 | 5,46E-05 |
| 215034_s_at | TM4SF1 | 23986 | 126,8 | 356,6 | 0,04734 | 5,61E-05 |
| 203685_at | BCL2 | 12949 | 130,2 | 320,1 | 0,04942 | 5,95E-05 |
